# MAP4K4 regulates biomechanical forces at adherens junctions and focal adhesions to promote collective cell migration

**DOI:** 10.1101/2022.11.22.517545

**Authors:** Lara Elis Alberici Delsin, Cédric Plutoni, Anna Clouvel, Sarah Keil, Léa Marpeaux, Lina Elouassouli, Adele Khavari, Allen Ehrlicher, Gregory Emery

## Abstract

Collective cell migration is important for normal development and tissue homeostasis, but can also promote cancer metastasis. To migrate collectively, cells need to coordinate their protrusion formation, rear retraction, adhesion sites dynamics, as well as forces generation and transmission. Nevertheless, the regulatory mechanisms coordinating these processes remain elusive. Using the A431 carcinoma cell line, we identify the kinase MAP4K4 as a central regulator of collective migration. We show that MAP4K4 inactivation blocks the migration of clusters while its overexpression decreases cluster cohesion. MAP4K4 regulates protrusion and retraction dynamics, remodels the actomyosin cytoskeleton, and controls the stability of both cell-cell and cell substrate adhesion. MAP4K4 promotes focal adhesion disassembly through the phosphorylation of Moesin, an actin and plasma membrane cross-linker, but disassembles adherens junctions through a Moesin-independent mechanism. By analyzing traction and intercellular forces, we found that the stabilization of adhesion sites in MAP4K4 loss of function leads to a tensional disequilibrium throughout the cell cluster, increasing the traction forces exerted onto the substrate and the tension loading at the cell-cell adhesions. Together, our results indicates that MAP4K4 activity is a key regulator of biomechanical forces at adhesion sites, promoting collective migration.

## INTRODUCTION

Collective cell migration is a highly coordinated process important for development, tissue homeostasis and wound healing. It can take several forms, since cells can migrate as sheets, streams, or clusters [1-3]. Collective cell migration can also occur in pathology, as during cancer metastasis. Increasing evidence has demonstrated that cancer cells migrating collectively are more efficient at forming metastases compared to individualized cells, as cell cluster are better at invading tissues and surviving in a new environment [4-6]. Moreover, patients presenting circulating tumor cell clusters have worse survival rates [7].

To migrate collectively, cells interact with their environment, frequently the extracellular matrix, and their neighbor migrating cells. To interact with their environment, cells form focal adhesions, large complexes of proteins that bridge the cell cytoskeleton to the extracellular matrix proteins, mainly through transmembrane proteins of the integrin family [8, 9]. The contact with the neighbor cells is mediated by cell-cell junctions, which connect the actomyosin cytoskeleton between two or more cells through cadherins and catenins [2, 10].

Both focal adhesions and adherens junctions are mechanosensitive platforms, where cells can apply, sense, and transmit forces. Forces are generated by the activation of myosin II, which binds to actin filaments, promoting cytoskeleton contraction. These forces can be transmitted through mechanosensitive protein-protein interactions at focal adhesions to generate traction forces, or at adherens junctions to promote intercellular stresses [10-12].

During collective migration, the adhesion sites, as well as the cytoskeleton structure, are constantly remodeled, with a net force that causes the cells to move. To metastasize collectively, neighboring cancerous cells need to coordinate the dynamics of cell-cell adhesions, promoting detachment from the primary tumor while retaining cluster cohesion [6]. Moreover, by remodeling their adhesions with the substrate and surrounding cells, cells can regulate the generation of traction forces and the transmission of stresses throughout the group [10]. How this communication mechanism is regulated is not entirely understood, and the central proteins that coordinate this process still need to be identified.

In this context, the Mitogen-Activated Kinase Kinase Kinase Kinase 4 (MAP4K4) influences collective cell migration in different model systems. MAP4K4 is a serine/threonine protein kinase of the Ste-20 family that has been implicated in the regulation of several signaling pathways. Therefore, MAP4K4 deregulation associates to different pathologies, including cancer [13, 14]. The Drosophila orthologue of MAP4K4, Misshapen, was highlighted as key for the coordination of protrusion extension and rear retraction during border cells cluster migration [15]. It was also shown to regulate focal adhesion dynamics in both Drosophila and mammalian cells, driving follicle epithelial cell migration during morphogenesis [16] and vascularization during mouse embryogenesis [17], respectively. Specifically, MAP4K4 induces integrin recycling, and different molecular models for this function have been proposed [17-19]. Importantly, MAP4K4 is overexpressed in several solid tumors and frequently associated with a poor survival rate [13, 20-22]. Increasing evidence places MAP4K4 as a pro-metastatic regulator, inducing cancer cell migration [13]. However, the role of MAP4K4 in collective migration of cancer cells has not been addressed.

Here we investigate the role of MAP4K4 in the regulation of the collective migration behavior of cancer cells, using the squamous epidermoid carcinoma cell line A431 as a model for cluster migration. We show that MAP4K4 is required for cluster migration through the regulation of protrusion and retraction dynamics. Specifically, MAP4K4 depletion stabilizes both the actomyosin cytoskeleton and focal adhesions, which results in higher traction forces at the substrate. We further report that higher forces are also applied on adherens junctions, and that intercellular stresses are regulated by MAP4K4. Interestingly, we found that MAP4K4 is recruited to adherens junctions where it promotes junction disassembly when overexpressed and, consequently, reduces force transmission. Overall, our work shows that MAP4K4 coordinates the generation and transmission of forces during collective cell migration, by regulating the stability of adhesion sites.

## RESULTS

### MAP4K4 is required for the collective migration of carcinoma cells through protrusion dynamics

When cultured at a low density, A431 cells form clusters of 6 to 15 cells that migrate collectively as a cohesive entity (see Material and Methods). Due to this property, A431 cell line was the primary model used for our study. To determine whether MAP4K4 is required for A431 cluster migration, we used CRISPR-Cas9 and two independent guides (sgRNA) to generate MAP4K4 knocked-out cells (MAP4K4 KO), or a non-target sgRNA sequence as control (sgNT) (Fig. 1a). Migrating cell clusters grown on collagen-matrigel gels were tracked over 5h. MAP4K4 KO reduced the instantaneous migration speed of clusters when compared to control sgNT (Fig. 1b, c). Moreover, treating cells with GNE-495, a specific MAP4K4 kinase inhibitor [23], reduced the migration speed in a dose dependent manner (Fig. 1d, e, Extended Data Fig. 1a), showing that the function of MAP4K4 in collective cell migration depends on its kinase activity. To gain insights into the processes regulated by MAP4K4, we tracked the margin of control and MAP4K4-inhibited cell clusters. We found that the displacement of the periphery was reduced after MAP4K4 inhibition (Fig. 1f, g), meaning that protrusions and cell retractions were less dynamic. Accordingly, the speed of both cellular extensions and retractions decreased (Fig. 1h, i, Extended Data Fig. 1b, c). This suggests that MAP4K4 regulates migration by promoting the dynamics of protrusion extensions and retractions across the cluster.

**Fig. 1.**
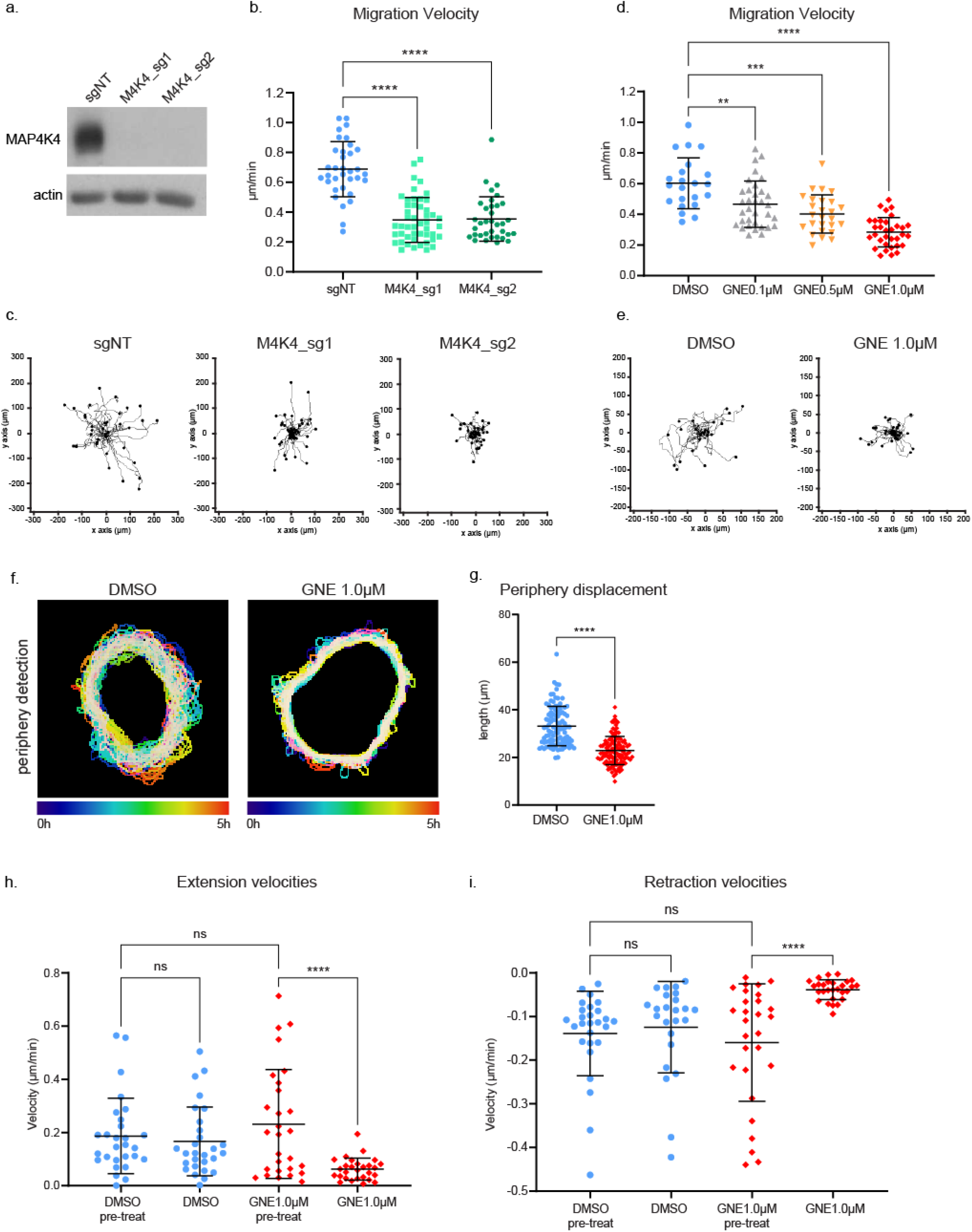
CRISPR/Cas9- or GNE495-mediated MAP4K4 inhibition impair A431 cluster migration. (a) representative immunoblotting of MAP4K4 and actin using lysates of A431 control cells (sgNT), or A431 cells KO for MAP4K4 with two independent sgRNA (M4K4_sg1, M4K4_sg2). (b) mean velocity of A431 clusters control or KO for MAP4K4, tracked over 5h of migration. (c) rose plots showing the cluster migration trajectory of A431 clusters control or KO for MAP4K4. (d) mean velocity of A431 clusters treated with DMSO or GNE-495 at different doses (0.1μM, 0.5μM or 1.0μM), over 5h of treatment. (e) rose plots showing the cluster migration trajectory of A431 clusters treated with DMSO or GNE-495 at 1.0μM. Number of clusters analyzed (sgNT: 34, M4K4_sg1: 48, M4K4_sg2: 35, DMSO: 22, GNE 0.1μM: 34, GNE 0.5μM: 26, GNE 1.0μM: 33), from 3 independent experiments. (f) projection of the cluster periphery automated detection, over 5h migration, color coded by time. Cells were treated with DMSO or GNE-495 at 1.0μM and the different timepoints were centered. (g) mean length of the periphery displacement of the clusters, from 0h to 15h of treatment with DMSO or GNE-495 at 1.0μM, calculated as described in the methods section. (h, i) Mean velocity extension (h) or retraction (i) events at the periphery of the clusters before or after treatment with DMSO or GNE-495 at 1.0μM, over 5h of treatment. Number of clusters analyzed (DMSO: 28, GNE 0.1μM: 26, GNE 0.5μM: 26, GNE 1.0μM: 28) from 3 independent experiments. Data in (b), (d), (h) and (i) were tested using Kruskal-Wallis and plotted as mean±s.d. Data in (g) was tested using Mann-Whitney and plotted as mean±s.d. \**p*<0.05, \*\**p*<0.01, \*\*\**p*<0.001, \*\*\*\**p*<0.0001.

To investigate how MAP4K4 regulates the dynamics of cell protrusion, we examined the actin cytoskeleton. Control clusters presented both protruding and retracting cells at their periphery, characterized respectively by apparent F-actin “arches” at the protrusion base (arrows) or retraction fibers (arrowheads) (Fig. 2a). On the other hand, both MAP4K4 KO and GNE-495 treated cells (from now on referenced as MAP4K4 loss of function – LOF) presented only cells with large, lamellipodia-like protrusions at the cluster periphery (Fig. 2b, c). Consequently, the morphology of the cluster was more circular (Fig. 2d). Similar morphological changes of MAP4K4 KO or GNE-495 treatment were observed when cells were treated with two other MAP4K4 inhibitors DMX-5804 [24] and PF-06260933 [25] (Extended Data Fig. 2a).

**Fig. 2.**
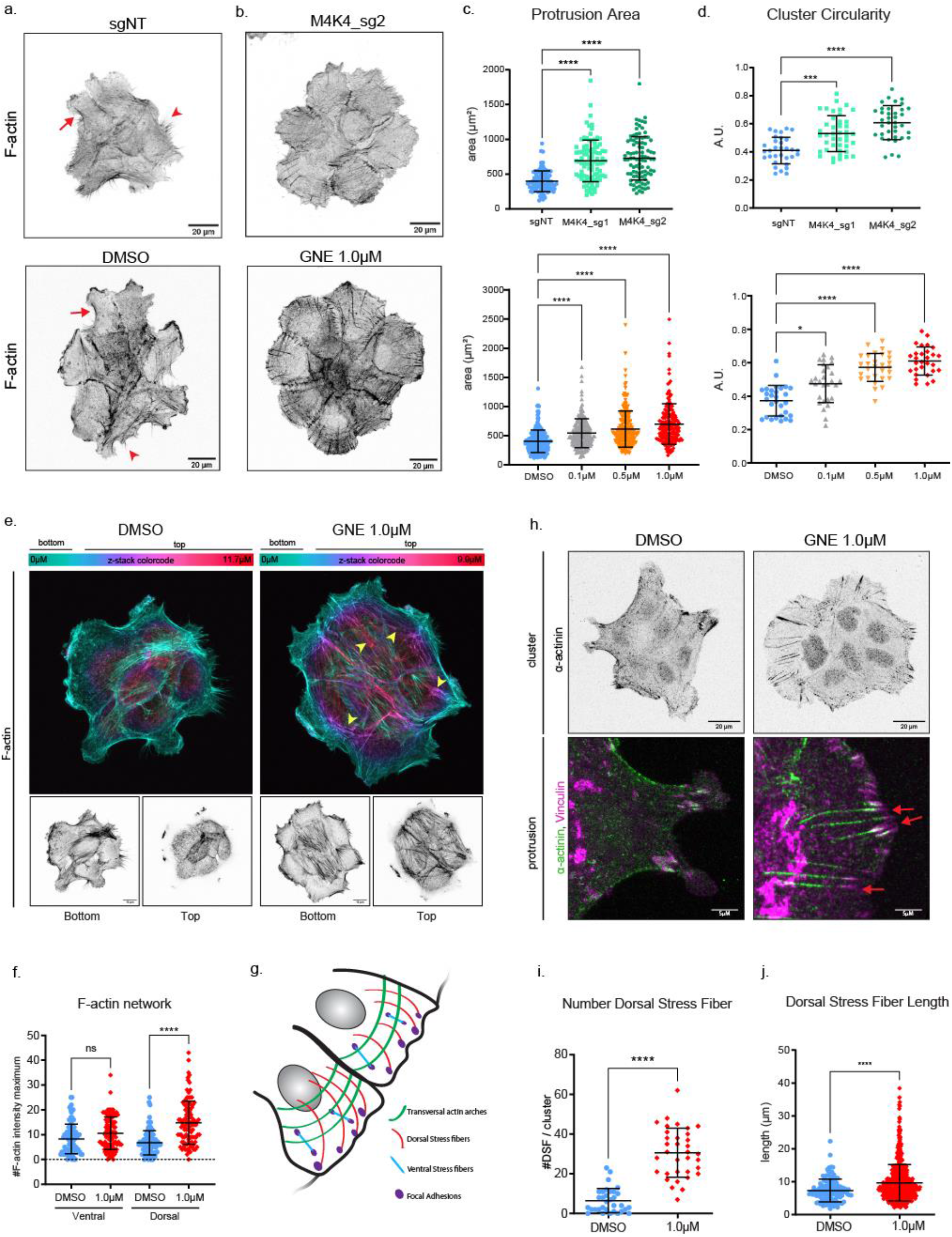
MAP4K4 LOF stabilizes protrusion and increases the F-actin bundles. (a) z-scan projection of representative confocal images of F-actin stained A431 clusters, showing the differences in the actin cytoskeleton organization and in the morphology of clusters control (sgNT) or KO for MAP4K4 (M4K4_sg2), or (b) clusters treated with DMSO or GNE-495 at 1.0μM for 24h. Arrows represent the actin arches at protrusion bases and arrowheads indicate retraction fibers. (c) protrusion area of control/MAP4K4 KO cells or DMSO/GNE-495 treated cells with indicated doses. At least 5 clusters per experiment, 3 protrusions per cluster from 3 independent experiments were analyzed. (d) circularity of control/MAP4K4 KO cell clusters, or clusters treated with DMSO or GNE-495 at indicated doses. At least 25 clusters from 3 independent experiments were analyzed. (e) images of representative clusters treated with DMSO or GNE-495 at 1.0μM stained for F-actin. Upper images are color-coded by the position in the z-axis, showing accumulation of thick F-actin at the dorsal part of a cluster treated with GNE-495, indicated by the increase of filaments in purple and magenta (arrowheads). Lower panel show projection of ventral z-positions (bottom) versus dorsal z-positions (top). (f) Relative abundance of F-actin bundles at ventral or dorsal part of clusters treated with DMSO or GNE-495 at 1.0μM, calculated as described in the methods section. At least 25 clusters from 3 independent experiments were analyzed. (g) Schematic representation of the different types of stress fibers (h) upper panel show confocal z-scan projection of representative A431 clusters treated with DMSO or GNE-495 at 1.0μM and stained for α-actinin. Bottom panel show a crop of the previous image showing cell protrusion stained for α-actinin (green) and vinculin (magenta), from clusters treated with DMSO or GNE-495 at 1.0μM. Arrows indicate F-actin fibers enriched in α-actinin that elongates from focal adhesions towards the dorsal part of the cluster, the so-called dorsal stress fibers. (i) number of dorsal stress fibers per cluster treated with DMSO or GNE-495 at 1.0μM. At least 30 clusters from 3 independent experiments were analyzed. (j) length of dorsal stress fibers of clusters treated with DMSO or GNE-495 at 1.0μM. All the dorsal stress fibers of at least 8 clusters per experiment from 3 independent experiments were measured. All the data is presented as mean±s.d. and tested by Kruskal-Wallis (\**p*<0.05, \*\**p*<0.01, \*\*\**p*<0.001, \*\*\*\**p*<0.0001).

Notably, MAP4K4 LOF induces the formation of a thick F-actin network, accumulating slightly in the ventral and mainly in dorsal regions of the cluster (Fig. 2e, f). These bundles are so-called stress fibers which support mechanical tension, helping cells to contract and to regulate their adhesion to the substrate [26]. Stress fibers can be classified as ventral or dorsal, as well as transversal arcs (Fig. 2g). Dorsal stress fibers extend from focal adhesions to the dorsal part of the cell and are enriched in α-actinin, which acts as a crosslinker to stabilize actin filament bundles. Dorsal stress fibers support the highly contractile transversal arcs, coupling the actomyosin machinery to focal adhesions [27, 28]. Immunostaining of α-actinin showed that MAP4K4 LOF induced the accumulation and elongation of dorsal stress fibers originating from focal adhesions (Fig. 2h-j). Those results indicate that MAP4K4 inhibition stabilizes the actin cytoskeleton at cell protrusions, reducing protrusion dynamics and slowing cell migration.

### MAP4K4 increases focal adhesion dynamics through Moesin phosphorylation

Previous work has shown that MAP4K4 regulates focal adhesion disassembly in different cell types [17-19]. To test if MAP4K4 is regulating focal adhesions on A431 carcinoma cells, we performed live imaging of the focal adhesion component paxillin fused to GFP. MAP4K4 inhibition decreased focal adhesion dynamics, as shown by the highly stable GFP enriched focal points over time (Fig. 3a, arrowheads, and Video 1).

**Fig. 3.**
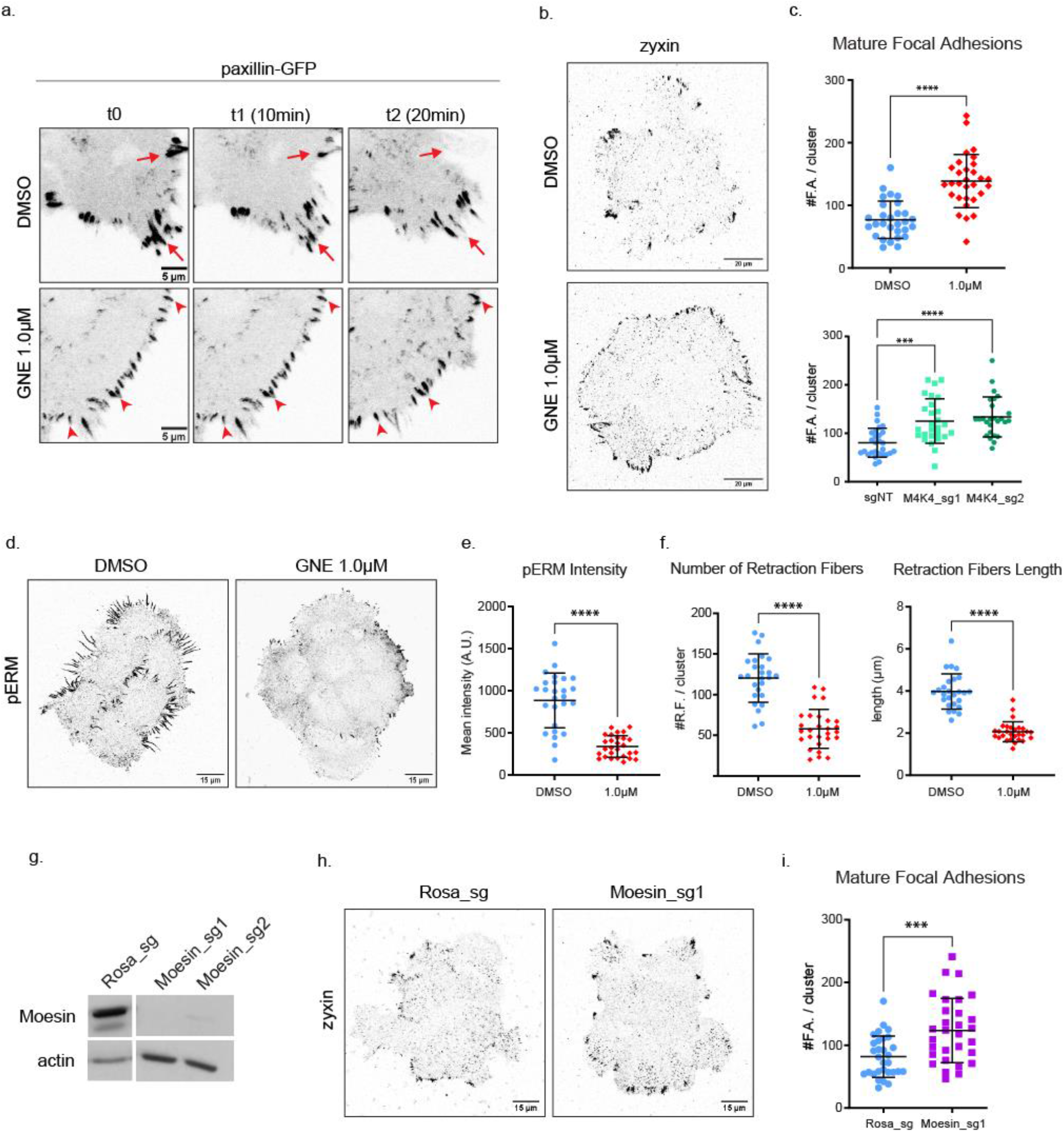
MAP4K4 LOF stabilizes focal adhesions and decreases Moesin phosphorylation. (a) confocal images of cell edges, acquired during a time-lapse of A431 cells expressing Paxillin-eGFP, treated with DMSO or GNE-495 at 1.0μM. Arrows indicate examples of focal adhesions that are disassembling over time, arrowheads indicate stable focal adhesions. (b) representative confocal images of A431 clusters treated with DMSO or GNE-495 at 1.0μM stained for zyxin, a mature focal adhesion marker. (c) number of zyxin-positive focal adhesions on clusters control (sgNT) or KO for MAP4K4, or treated with DMSO or GNE-495 at 1.0μM. At least 8 clusters per experiment, from 3 independent experiments were analyzed. (d) representative confocal images of pERM staining, at the substrate z focal plane, from clusters treated with DMSO or GNE-495 at 1.0μM. (e) quantification of mean intensity of pERM from clusters treated with DMSO or GNE-495 at 1.0μM. At least 25 clusters from 3 independent experiments were analyzed. (f) number and length of retraction fibers of clusters treated with DMSO or GNE-495 at 1.0μM. At least 25 clusters from 3 independent experiments were analyzed. (g) representative immunoblotting of Moesin and actin from lysates of A431 cells control (Rosa_sg) or KO for Moesin using 2 independent sgRNA sequences (Moesin_sg1, Moesin_sg2). (h) representative confocal images of control or Moesin KO clusters stained for zyxin. (h) number of zyxin-positive focal adhesions in control or Moesin KO clusters. (i). All the data is presented as mean±s.d. and tested by Mann-Whitney (\**p*<0.05, \*\**p*<0.01, \*\*\**p*<0.001, \*\*\*\**p*<0.0001).

Focal adhesions are formed as nascent adhesions at the front of the lamellipodium, and the subsequent recruitment of structural and signaling components induce their maturation while moving rearwards to the lamella. Mature focal adhesions bind to the cytoskeleton stress fibers through the actin binding proteins α-actinin, zyxin and VASP, and generate traction forces at their distal tip [8, 29, 30]. Since MAP4K4 LOF stabilizes focal adhesions and increases the number and length of stress fibers, we hypothesized that the number of mature focal adhesions would increase. Immunostaining of zyxin demonstrated that the number of mature adhesions indeed increased in MAP4K4 LOF cells (Fig. 3b, c, Extended data Fig. 2b).

In endothelial cells, MAP4K4 disassembles focal adhesions through the local phosphorylation of Moesin [17]. Moesin is a member of the ERM (Ezrin, Radixin, Moesin) family of proteins that link cortical actin to the plasma membrane [31]. Phosphorylation of Moesin by MAP4K4 ultimately induces integrin inactivation and focal adhesion disassembly [17]. To test if a similar mechanism is at play in A431 cells, we monitored phosphorylated ERM (pERM) using a phospho-specific antibody that recognizes a conserved phosphosite in all ERM proteins [32]. We observed that MAP4K4 inhibition decreases pERM intensity at the focal plane containing the focal adhesions, suggesting that ERM proteins are phosphorylated by MAP4K4 at the substrate focal plane (Fig. 3d, e). Concomitantly, we observed a reduction in the number and length of retraction fibers, which are ERM enriched structures that are formed when cells retract (Fig. 3f).

To determine if MAP4K4 LOF stabilizes focal adhesion through the regulation of a specific ERM protein, we generated Ezrin, Radixin and Moesin KO cells by CRISPR-Cas9, using two independent guide sequences (sgRNA) for each family member. The Moesin KO cells presented an increase in zyxin-enriched mature focal adhesions (Fig. 3g-i), while Ezrin and Radixin KOs showed no difference when compared to control (data not shown). Altogether, our data suggests that MAP4K4 phosphorylates Moesin to regulate the dynamics of focal adhesion also in A431 epidermoid cells.

### MAP4K4 decreases generation of traction force during CCM

Extensive literature shows that myosin II activity is required at cell protrusions for focal adhesion maturation, stress fiber elongation, and increasing traction forces [8, 28, 33, 34]. Because MAP4K4 LOF induced accumulation of both mature focal adhesion (Fig. 3) and stress fibers (Fig. 2), we investigated if it would also affect myosin II-mediated contractility. To test if MAP4K4 LOF would impact the activation of myosin II, we measured the phosphorylation status of the myosin light chain 2 (MLC2) by western blot using a phospho-specific antibody. We observed no detectable difference on myosin phosphorylation levels after the inhibition of MAP4K4 (Fig. 4a, b).

**Fig. 4.**
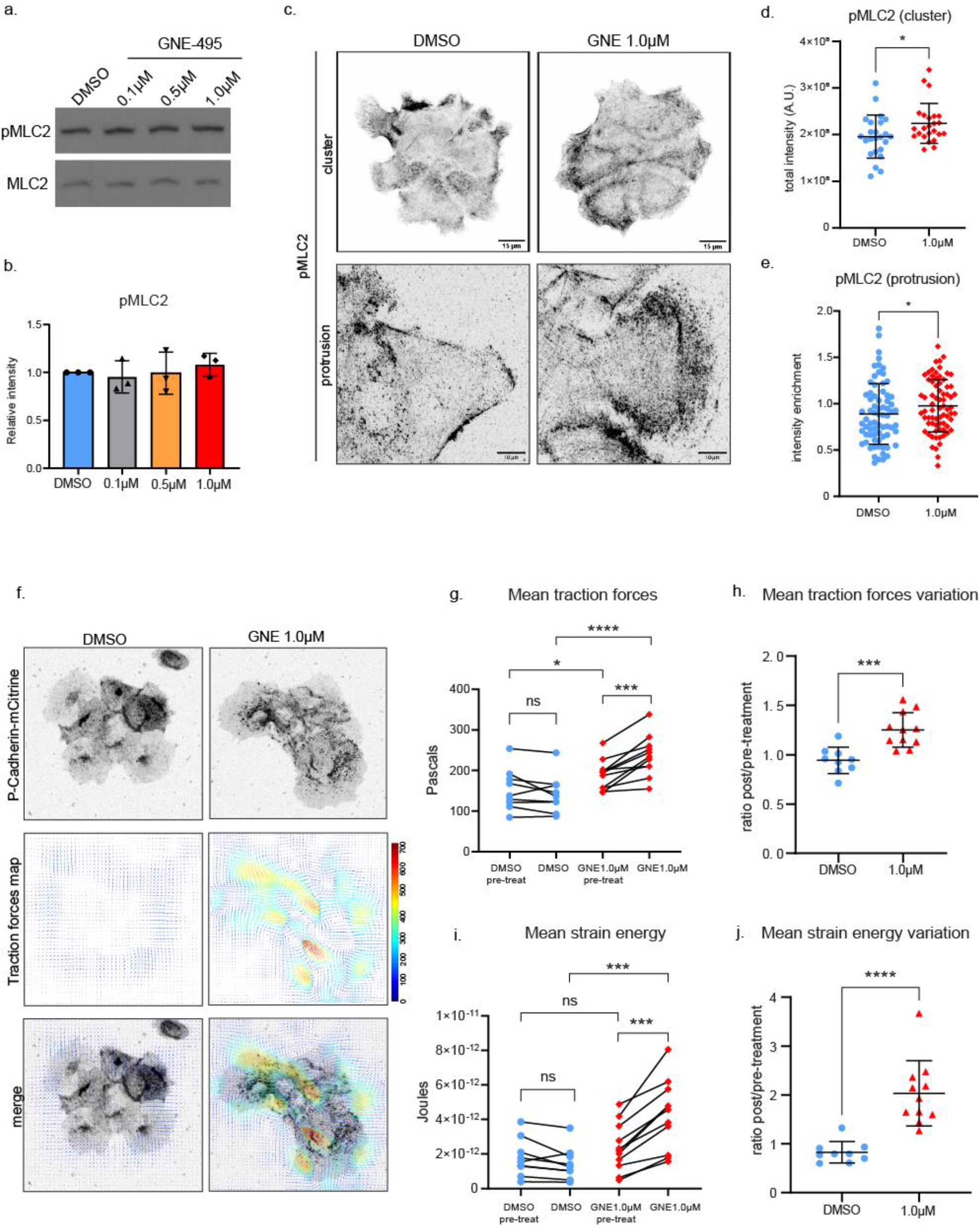
MAP4K4 LOF increases contractility at protrusion and cluster traction forces. (a) representative immunoblotting of pMLC2 and MLC2 using lysates of A431 cells treated with DMSO or indicated doses of GNE-495. (b) ratio of pMLC2/MLC2 intensities from 3 independent experiments of A431 clusters treated with the indicated doses of GNE-495, relative to DMSO. (c) z-scan projections of representative confocal images of pMLC2 stained A431 clusters (upper panel) or cell protrusion (lower panel), showing the differences in pMLC2 accumulation in DMSO or after GNE-495 treatment at 1.0μM. (d) quantification of total pMLC2 intensity in DMSO or after GNE-495 treatment. At least 24 clusters from 3 independent experiments were analyzed. (e) ratio of the mean intensity of pMLC2 at protrusions over the mean intensity of pMLC2 at entire cluster. At least 3 protrusion per cluster, from 8 clusters per experiments of 3 independent experiments were analyzed. (f) confocal images acquired during time-lapse of A431 cells expressing P-Cadherin-mCitrine, treated with DMSO or GNE-495 at 1.0μM. Representative images of traction maps and overlay of traction maps and fluorescent images. (g) mean traction forces of individual clusters before or after treatment with DMSO or GNE-495 at 1.0μM (average of 3 timepoints before treatment or between 4h30-5h of treatment). (h) ratio of mean traction forces of individual clusters treated with DMSO or GNE-495 over the paired cluster before treatment. (i) mean strain energy of individual clusters before or after treatment with DMSO or GNE-495 at 1.0μM (average of 3 timepoints before treatment or between 4h30-5h of treatment). (j) ratio of mean strain energy of individual clusters treated with DMSO or GNE-495 over the paired cluster before treatment. At least 9 clusters from 3 independent experiments were analyzed. Data on (b), (d), (e), (h) and (j) are represented as mean±s.d. and tested by Mann-Whitney. Unpaired analysis on (g) and (i) was done by Mann-Whitney test, while paired analysis was done using Wilcoxon test (ns: nonsignificant, \**p*<0.05, \*\**p*<0.01, \*\*\**p*<0.001, \*\*\*\**p*<0.0001).

We next investigated whether the localization of active myosin II is affected. In contrast to the western blot analysis, quantification of the immunostaining of the phosphorylated active form of MLC2 (pMLC2) revealed a small, but significant increase in the total levels of active myosin II in MAP4K4 LOF (Fig. 4c, d). More importantly, we observed an enrichment of active myosin at cell protrusion (Fig. 4c, e), indicating that MAP4K4 LOF induces a relocation of active myosin to the lamella, where it may contribute to the maturation of the focal adhesion and elongation of stress fibers [8, 28, 35]. Such alterations of the organization of protrusions might cause an increase in the traction forces exerted on the substrate.

To determine if the inhibition of MAP4K4 has an impact on the forces applied to the substrate, we used Traction Force Microscopy (TFM) [36, 37]. Treatment of A431 clusters with GNE-495 induced higher mean traction forces and mean strain energy when compared to treatment with DMSO (Fig. 4f-j). Therefore, when MAP4K4 is inactive, cells exert higher traction forces on the substrate, showing that MAP4K4 activity influences tension at cell-substrate interface.

Altogether, those results indicate that the absence of MAP4K4 triggers a mechanosensitive-like response in the cell. This response consists of 1) the stabilization of focal adhesions, 2) the maturation of those structures by binding and elongating α-actinin enriched stress fibers, 3) the recruitment of active myosin to the protrusion lamella and 4) the increase in the traction forces exerted on the substrate. However, the temporal and casual relationship between these effects need to be further investigated.

### MAP4K4 decreases tension at adherens junction

During collective cell migration, forces generated at the substrate can be transmitted to the neighbor cells through adherens junctions [11, 38-40]. Since MAP4K4 LOF induces an increase in traction forces, we asked whether it would also affect the adherens junctions and the transmission of those forces. Staining of the adherens junction marker p120 catenin revealed junction alterations in MAP4K4 LOF clusters. In control clusters, adherens junctions are mostly linear, while the junctions of MAP4K4 LOF cells present a tortuous morphology (Fig. 5a-d).

**Fig. 5.**
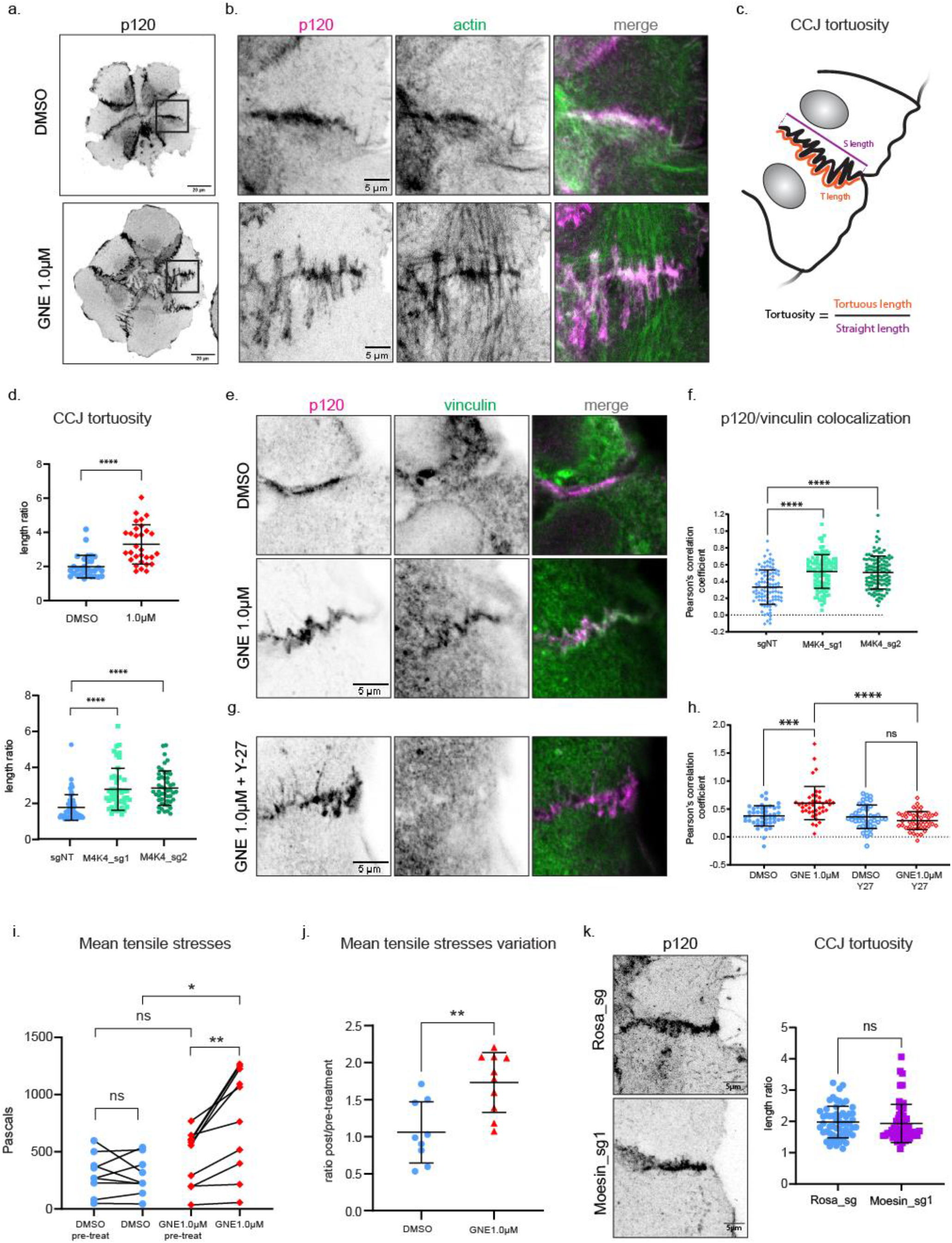
MAP4K4 LOF increases tension loading at cell-cell junctions. (a) z-scan projection of representative confocal images of p120-catenin stained A431 clusters, showing the differences in the cell-cell junction morphology of clusters treated with DMSO or GNE-495 at 1.0μM (b) crops of the previous images, indicating parallel or perpendicular F-actin organization at the adherens junction. (c) Schematic representation of cell-cell junction tortuosity index calculation (d) cell-cell junction tortuosity index for A431 clusters control (sgNT) or KO for MAP4K4, or treated with DMSO or GNE-495 at 1.0μM. At least 3 junctions of 5 different clusters per experiment, from 3 independent experiments were analyzed. (e) representative confocal images of cell-cell junctions, showing vinculin accumulation at junctions of clusters treated with DMSO or GNE-495 at 1.0μM. colocalization between p120 and vinculin intensities for clusters control (sgNT) or KO for MAP4K4, calculated by the Pearson’s correlation coefficient. At least 3 junctions per cluster, from at least 8 clusters per experiment, from 3 independent experiments were analyzed. (g) representative confocal images of cell-cell junctions, showing loss of vinculin accumulation at junctions when GNE-495 treated clusters for 24h were exposed to the ROCK inhibitor Y-27632 during 15min. (h) colocalization between p120 and vinculin in clusters treated with DMSO or GNE-495 at 1.0μM alone or in combination with Y-27632 (2.5μM), calculated by the Pearson’s correlation coefficient. At least 3 junctions per cluster, from at least 8 cluster per experiment, from 3 independent experiments were analyzed. (i) BISM analysis yielded the intercellular stresses, summarized here by tensile stresses. Mean tensile stresses were plotted as individual clusters before or after treatment with DMSO or GNE-495 at 1.0μM (average of 3 timepoints before treatment or between 4h30-5h of treatment). (j) ratio of mean tensile stresses of individual clusters treated with DMSO or GNE-495 over the paired cluster before treatment. At least 9 clusters from 3 independent experiments were analyzed. (k) representative confocal z-scan projection of p120 stained cells, control or KO for Moesin, showing cell-cell junction morphology. Graph indicates tortuosity index of cell-cell junction on control or Moesin KO cells. At least 3 junctions of 5 different clusters per experiment, from 3 independent experiments were analyzed. Data on (d), (j) and (k) are represented as mean±s.d. and tested by Mann-Whitney. Data on (f) and (h) are represented as mean±s.d. and tested by Kruskal-Wallis. Unpaired data on (i) was analyzed by Mann-Whitney test, while paired data was analyzed using Wilcoxon test (ns: nonsignificant, \**p*<0.05, \*\**p*<0.01, \*\*\**p*<0.001, \*\*\*\**p*<0.0001).

A similar tortuous adherens junction shape was also reported in endothelial cells. They can appear after chemically induced contractility and are perpendicularly bound to stress fibers of neighbor cells [41, 42], as seen in MAP4K4 LOF cells (Fig. 5b). Based on both, the increase on traction forces and the reshaping of adherens junction, we decided to further investigate whether MAP4K4 regulates tension loading on cell-cell adhesions.

To test that, we immunostained vinculin, a protein recruited to adherens junctions under tension, downstream of opening of the mechanosensitive protein α-catenin, whose density increases with tension [43-45] MAP4K4 LOF clusters accumulate vinculin at the adherens junctions compared to control (Fig. 5e, f), suggesting that the adherens junctions are under higher tension when MAP4K4 is inactive. To understand if this phenotype is dependent on actomyosin contractility, we combine the GNE-495 treatment with a contractility inhibitor, the Rho kinase (ROCK) inhibitor (Y-27632). The addition of Y-27632 abrogates the vinculin recruitment at adherens junction induced by MAP4K4 LOF, showing that this recruitment requires myosin induced contractility (Fig. 5g, h). Those results suggest that MAP4K4 decreases contractility and tension loading also at the adherens junctions.

Moreover, we used our TFM data to estimate the intercellular stresses field in A431 clusters, by Bayesian Inversion Stress Microscopy analysis (BISM). We found that intercellular tensile stresses are substantially increased after inhibition of MAP4K4 (Fig. 5i, j), suggesting that those tensile stresses are sensed and transmitted across the entire cluster. Together, those results indicate that MAP4K4 decreases tension loading at adherens junctions and regulates the magnitude of the intercellular stresses.

Aiming at deciphering the molecular mechanism by which MAP4K4 regulates tension loading at the adherens junction, we started by testing whether Moesin phosphorylation was also involved with the role of MAP4K4 on cell-cell junction. Thus, we stained pERM by immunofluorescence and measured the signal intensity at the adherens junction focal plane in control or MAP4K4 inhibited clusters. We did not observe any change in pERM intensity at that focal plane in MAP4K4 LOF cells (Extended Data Fig. 3a, b). Moreover, Moesin KO clusters do not present tortuous junctions (Fig. 5k), showing that the loss of Moesin activity is not sufficient to induce the MAP4K4 phenotype at adherens junction. Altogether our data suggest that MAP4K4 acts on a different substrate at adherens junction.

Because Moesin KO increases the number of mature focal adhesions (Fig 3h, i), but does not make cell-cell junctions more tortuous (Fig. 5k), we inferred that the effect of MAP4K4 KO at cell-cell junctions is not an indirect effect of focal adhesion stabilization. Hence, MAP4K4 might directly regulate forces at adherens junctions.

### MAP4K4 localizes at adherens junctions and regulates their disassembly

To examine if MAP4K4 has a direct effect on adherens junctions, we investigated its localization with an eGFP fusion to MAP4K4. We found that eGFP-MAP4K4 localizes at adherens junctions (Fig. 6a), supporting our hypothesis that MAP4K4 directly regulates tension at junctions. Interestingly, we frequently observed that MAP4K4 accumulates at the interface of cells that are in close proximity, but not presenting continuous cell-cell junctions, as we observe many gaps between those cells (Fig. 6b). Time-lapse recordings show that shortly after the formation of cell-cell contacts (red arrows), eGFP-MAP4K4 accumulates at the cell-cell interface (cyan arrows). Moreover, MAP4K4 accumulation at those newly formed cell-cell junctions is followed by the disassembly of those adhesion sites (Video 2). Based on these observations, we can hypothesize that recruitment of MAP4K4 at cell-cell junctions promotes its disassembling. During the time-lapse recording and on fixed samples, we also observe that MAP4K4 localizes at retraction fibers and cell rear in both single cells and cells in clusters (yellow arrowheads on Video 2, Extended data Fig. 4a, b). Previous work showed that MAP4K4 is present at retraction fibers in endothelial cells [17]. From our data, we can conclude that 1) the localization of MAP4K4 at restriction fibers is not cell type specific, 2) MAP4K4 is not restricted to retraction fibers as it is present near the retraction areas and at cell-cell junctions.

**Fig. 6.**
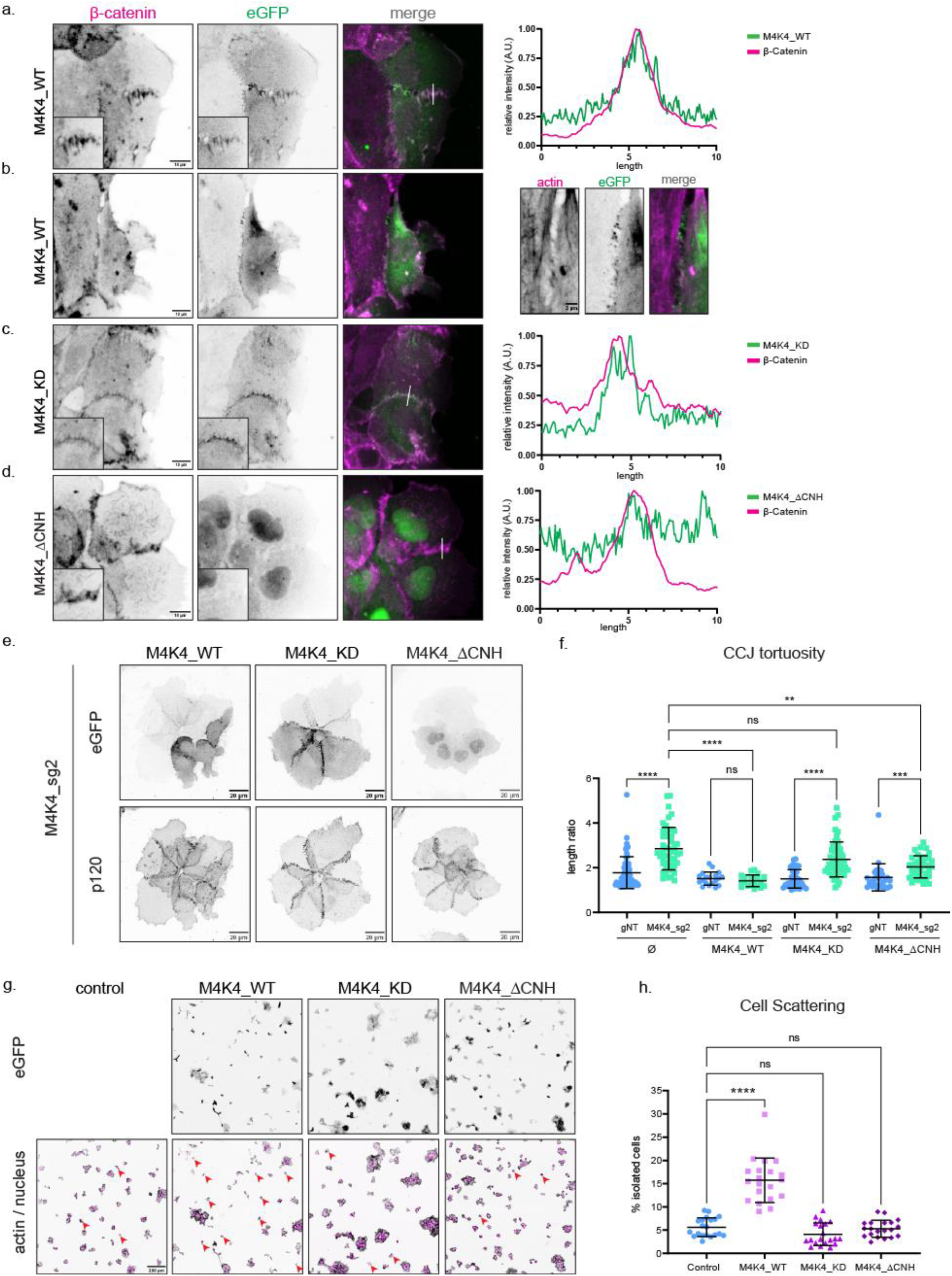
MAP4K4 localizes at cell-cell junction, in a CNH dependent manner, and induces cell scattering. (a) z-scan projection of representative confocal images of β-catenin stained A431 clusters, stably expressing eGFP-MAP4K4_WT. Line scan indicates the colocalization between β-catenin and MAP4K4. (b) z-scan projection of confocal image exemplifying the localization of MAP4K4_WT at spots where cells seem to detach from their neighbors. (c) z-scan projection of representative confocal images of β-catenin stained A431 clusters, stably expressing eGFP-MAP4K4 kinase dead (MAP4K4^D153N^) or (d) deleted for the CNH domain (MAP4K4^ΔCNH^). Line scans indicate the presence (for kinase dead) or absence (for CNH deletion) of MAP4K4 at cell-cell junctions. (e) z-scan projection of representative confocal images of A431 clusters KO for MAP4K4, expressing exogenous eGFP-MAP4K4 WT, KD or ΔCNH constructs with silent mutations on sg2 targeted region (see material and methods). The p120-catenin staining was used to visualize the cell-cell junction morphology. (f) cell-cell junction tortuosity index for A431 clusters control (sgNT) or KO MAP4K4 (M4K4_sg2), and stably expressing eGFP-MAP4K4 WT, KD or ΔCNH, resistant to sg_2. (g) representative confocal images of A431 cells and clusters stably expressing eGFP-MAP4K4 WT, KD or ΔCNH. Arrowheads indicate examples of isolated cells in each condition. (h) Percentage of cell scattering in A431 control, or overexpressing eGFP-MAP4K4 WT, KD or ΔCNH. Data on (f) and (h) are represented as mean±s.d. and tested by Kruskal-Wallis (ns: nonsignificant, \**p*<0.05, \*\**p*<0.01, \*\*\**p*<0.001, \*\*\*\**p*<0.0001).

To explore how the localization of MAP4K4 at adherens junctions is regulated, we generated mutant constructs. MAP4K4 is composed of a kinase domain at its N-terminus, followed by a coiled-coil and an unstructured region, and a CNH domain (citron homology domain) at its C-terminal [14]. We generated eGFP tagged constructs with a kinase-inactive mutant (MAP4K4^D153N^), as well as a C-terminal deletion of its CNH domain (MAP4K4^ΔCNH^) and explored their localization. We found that the recruitment of MAP4K4 at adherens junction is independent of its kinase activity, however, recruitment does require the CNH domain (Fig. 6c, d). The localization of MAP4K4 and its mutants are similar in monolayers of MDCK cells, epithelial cells derived from canine kidney (Extended Data Fig. 4c), suggesting that the localization of MAP4K4 at adherens junction is not unique to a single cell type.

We further tested the functionality of those constructs by performing rescue experiments in MAP4K4 KO cells. The wildtype form of MAP4K4 (MAP4K4^wt^) completely restores junction linearity in KO clusters. However, KO cells expressing the kinase-inactive mutant still present a higher tortuosity, while the expression of MAP4K4^ΔCNH^ induces only a partial rescue (Fig. 6e, f, Extended data Fig. 4d, e). Those results indicate that both the CNH domain and the kinase activity are necessary for the full function of MAP4K4 at adherens junctions. Moreover, the kinase-inactive construct corroborates that the effect observed upon GNE-495 treatment is specific to MAP4K4 kinase activity impairment.

Interestingly, we observed that MAP4K4^wt^ overexpression induces a higher percentage of individualized cells among the A431 clusters, which is not seen after kinase-inactive mutant or MAP4K4^ΔCNH^ overexpression (Fig. 6g, h, Extended data Fig. 4f). Together with its localization at the sites of remnant adhesions (Fig. 6b) and the disassembly of cell-cell contacts after MAP4K4 accumulation (Video 2), those results suggest that MAP4K4 regulates cell-cell junction disassembly and promotes cell scattering.

## DISCUSSION

In this work we used the epidermoid carcinoma cell line A431 as a model to study the role of the kinase MAP4K4 in regulating the collective migration of cancer cell clusters in vitro. We show that MAP4K4 is a central regulator of force generation and transmission during collective migration, acting specifically on the disassembly of cell-substrate and cell-cell adhesions. Previous work showed that MAP4K4 regulates the disassembly and recycling of focal adhesions, through different molecular mechanisms [17-19]. Here, we found that MAP4K4 induces focal adhesion turnover predominantly through Moesin phosphorylation in A431 carcinoma cells. Moreover, we show that when MAP4K4 is impaired there are more mature focal adhesions, leading to an increase in the number of stress fibers at cell protrusions, and a decrease in protrusion dynamics. Therefore, MAP4K4 inhibition not only prevents cells to retract, but reorganizes the cytoskeleton network inside protrusions. Furthermore, we reveal that this process relocates active myosin to those protrusions, locally increasing contractility and inducing cells to exert higher traction forces on the substrate (Fig. 7a). In the collective context, the presence of protrusions around the entire cluster, as observed in MAP4K4 LOF, impairs the cell-cell coordination mechanism and blocks migration [46]. Therefore, the stabilization of focal adhesion by MAP4K4 LOF initiates a cascade of cellular processes that culminates in the impairment of collective cell movement (Fig. 7b, c).

**Fig. 7.**
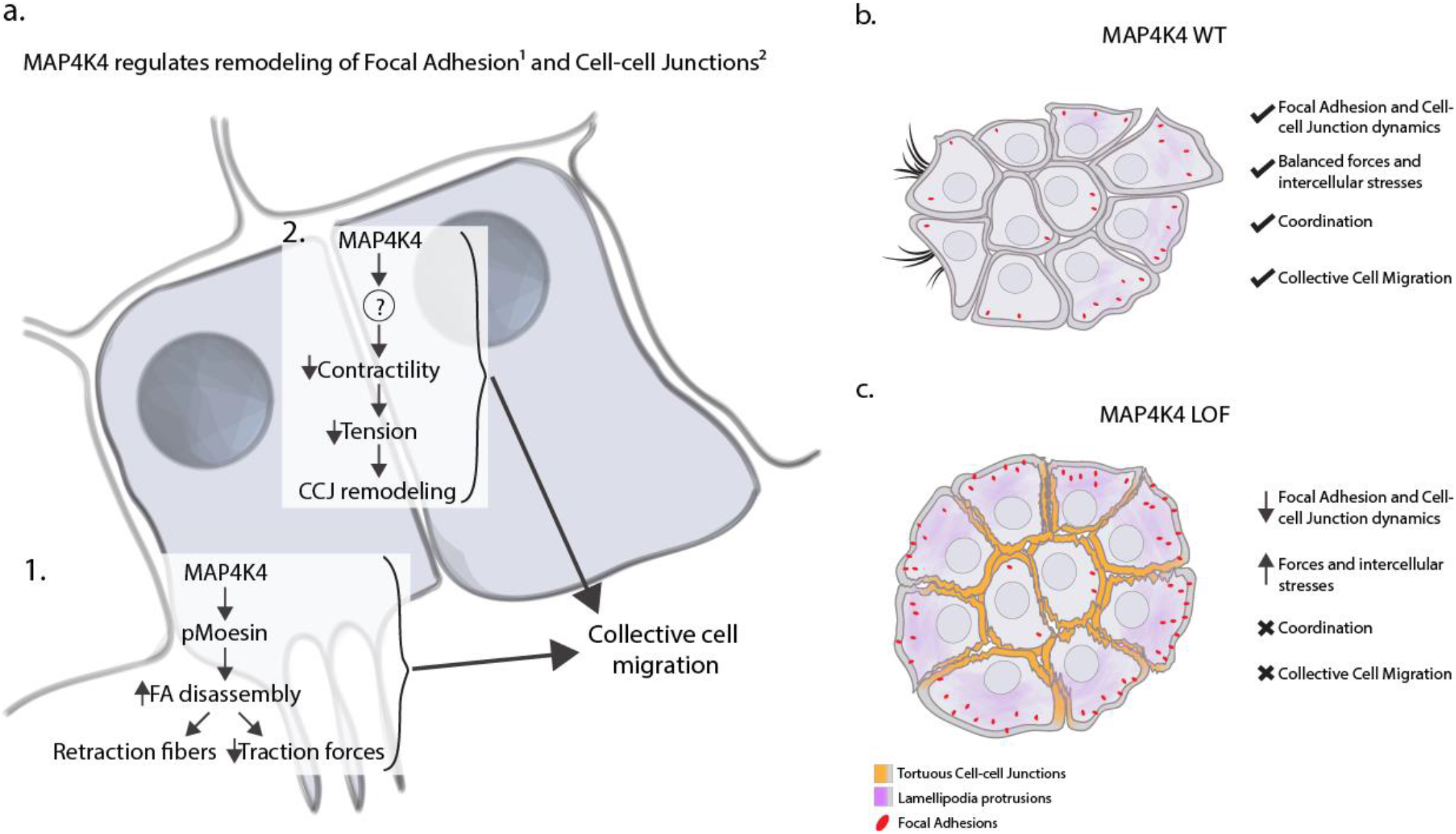
MAP4K4 acts on focal adhesion and adherens junctions to regulate collective cell migration. (a) Proposed model depicting how MAP4K4 regulates 1. focal adhesion disassembly and 2. cell-cell junction remodeling, allowing a tensional equilibrium and promoting collective cell migration. (b, c) summary of the main biomechanical processes necessary for cluster migration, which are deregulated by MAP4K4 loss of function.

We also report that MAP4K4 can be recruited to cell-cell adhesions, and this seems to decrease their stability, tension loading and transmission of forces (Fig. 7a-c). Previous work has suggested a role of MAP4K4 in cell-cell adhesion stability by studying endothelial cells’ permeability. In that context, MAP4K4 depletion increases the resistance of the endothelial barrier [47], which can indicate a tightening of cell-cell junctions. Here, we show that MAP4K4 LOF induces higher cell-cell junction tortuosity, which physically increases the adhesion area between the cells. We also show the recruitment of vinculin to cell-cell junctions in MAP4K4 LOF cells. Vinculin is recruited to the adherens junctions to help stabilizing the catenin-actin complex when strong pulling forces are exerted [43, 45, 48]. Therefore, accumulation of vinculin in MAP4K4 LOF indicates higher junction stabilization and increase of tension loading, as validated by our intercellular stress calculations. Conversely, we show that MAP4K4 overexpression induces adhesion disassembly and cell scattering. Those results reinforce the idea that MAP4K4 balances the adhesiveness and stability of adherens junctions under increased pulling forces.

Interestingly, Pannekoek and colleagues report that MAP4K4 acts downstream the small GTPase Rap2, which binds to the CNH domain of MAP4K4 [47, 49]. Here, we describe that the recruitment of MAP4K4 to adherens junction depends on its CNH domain. Therefore, it is appealing to hypothesize that MAP4K4 may be recruited to adherens junctions by Rap2 to decrease its stabilization upon mechanical stresses. Accordingly, Meng and collaborators found that MAP4K4 acts downstream of Rap2 activation in response to substrate stiffness and focal adhesion stabilization [50]. Therefore, it would be interesting to test if a similar mechanoresponse involving Rap2 and MAP4K4 is at play at adherens junctions.

Moreover, other proteins that contain a CNH domain are also known to bind active Rho GTPases, as RhoA, Rac1 and Cdc42 [51, 52]. Rho GTPases are master regulators of cell cytoskeleton and contractility [53, 54], as well as junction assembly and maintenance [53, 55, 56]. Therefore, MAP4K4 might also be recruited to adherens junctions through the binding of Rho GTPases.

Despite the importance of MAP4K4 for cancer progression, little is known about its downstream direct targets. At focal adhesions, MAP4K4 seems to act predominantly through Moesin phosphorylation. However, our data suggest that this is not the case at adherens junctions. Although there are few known direct substrates that could be at play at adherens junction, they do not seem to explain MAP4K4 LOF phenotype in A431 clusters. One of MAP4K4 targets is the actin nucleator Arp2 [57], however, it is unlikely that Arp2 is downstream of MAP4K4 at junctions. Arp2 is required for the maintenance of cell-cell junction and does not promote adhesion disassembly as MAP4K4 [58-60]. Another known MAP4K4 substrate is LATS1/2, which regulates the mechanosensitive Hippo signaling pathway. Although LATS is an interesting potential target of MAP4K4 in A431 cells, Meng and collaborators [61] showed that depletion of MAP4K4 alone is not sufficient to inactivate the Hippo transcriptional factor YAP. Therefore, it is unlikely that MAP4K4 regulates cell-cell adhesion through LATS, although some contribution of the Hippo pathway cannot be excluded.

Therefore, we think that MAP4K4 is acting directly at adherens junctions through a yet unidentified substrate. By exploring The Human Cell Map database [62], we found that MAP4K4 is predicted to localize at cell-cell junctions, consistent with our findings. Furthermore, MAP4K4 potentially interacts with several junctional proteins, including afadin, α-catenin and occludin. Future work will have to establish if MAP4K4 interacts physically or directly phosphorylates these proteins.

In conclusion, our findings show that MAP4K4 promotes the disassembly of both focal adhesions and adherens junctions. Focal adhesions and adherens junctions are indirectly connected through the cell cytoskeleton and several proteins are shared between the two structures [38, 63]. The mechanical crosstalk between them has been explored [11, 64] and they can present cooperating [39] or antagonistic [65] responses. Here, we report a functional mechanism where MAP4K4 activity is central for balancing traction force generation through disassembly of focal adhesions, while it is also required to decrease the intercellular stresses and tension loading at the adherens junctions. Our work indicates that MAP4K4 is a key regulator of force balance to promote the collective migration of carcinoma cells. By regulating MAP4K4 expression and/or activation levels, cancer cell clusters can modulate their level of cohesion and thus the nature of their collective migration properties. Therefore, our results highlight a potential explanation for MAP4K4 pro-metastatic behavior.

## MATERIALS AND METHODS

### Antibodies

For immunoblotting, the following primary antibodies were used: rabbit polyclonal anti-HGK (MAP4K4) at 1:1000 (Cell Signaling Technology, #3485), rabbit polyclonal anti-phospho-Myosin Light Chain 2 (Ser19) at 1:1000 (Cell Signaling Technology, #3671), rabbit polyclonal anti-Non-Muscle Myosin HC II-A at 1:1000 (Biolegend, #909802), mouse monoclonal anti-Actin at 1:10000 (Millipore (C4), MAB1501), rabbit polyclonal anti-Moesin at 1:1000 (Cell Signaling Technology (Q480), #3150)

And the following secondary antibodies: AffiniPur goat anti-rabbit (H+L) and goat anti-mouse (H+L) (Jackson Immunoresearch, 111-035-144 and 115-035-062, respectively, used at 1/10000).

For immunofluorescence, the following primary antibodies were used: rabbit polyclonal anti-phospho-Ezrin (Thr567)/Radixin (Thr564)/Moesin (Thr558) at 1:250 (Cell Signaling Technology, #3141), rabbit polyclonal anti-phospho-Myosin Light Chain 2 (Ser19) at 1:500 (Cell Signaling Technology, #3671), rabbit polyclonal anti-α-actinin at 1:400 (USBiological, A0761-2F), mouse monoclonal anti-zyxin at 1:200 (SantaCruz (2D1), sc-293448), rabbit polyclonal anti-p120 catenin at 1:200 (SantaCruz (H90), sc-13957), mouse monoclonal anti-β-catenin at 1:100 (BD Biosciences, 610153), mouse monoclonal anti-vinculin at 1:200 (Sigma, V9131).

And following secondary antibodies, at 1:1000 dilution: Alexa Fluor 488 goat anti-mouse IgG (Invitrogen, A11029), Alexa Fluor 488 goat anti-rabbit IgG (Invitrogen, A11008), Alexa Fluor 555 anti-mouse IgG (Cell Signaling Technology, 4499S), Alexa Fluor 555 anti-rabbit IgG (Cell Signaling Technology, 4413S).

To stain nuclei, we used DAPI at 1μg/ml (Sigma, D8417-10MG).

To stain F-actin, we used Alexa Fluor 488 phalloidin at 1:1000 (Invitrogen, A12379), Alexa Fluor 555 phalloidin at 1:750 (Invitrogen, A34055), Alexa Fluor 647 phalloidin at 1:50 (Invitrogen, A22287).

### Cell Culture and plasmid transfection

A431 (CRL-1555, ATCC), MDCK (CCL-34, ATCC) or HEK-293-T (CRL-3216) cells were cultured in DMEM (Sigma-Aldrich), supplemented with 10% Fetal Bovine Serum (Gibco) and 1% penicillin-streptomycin (Sigma-Aldrich) in 5% CO_2_ air humidified atmosphere at 37°C. For A431 cells, cluster confluence is reached by plating 1500 - 2000 cells/cm^2^ and keeping cells in culture for 3 days. Cells were sporadically tested for mycoplasma. A431 plasmid transfection was done by using Lipofectamine™ 3000 Transfection Reagent (Thermo Fisher Scientific) following the manufacture’s recommendations.

### Constructs, CRISPR and virus production

pGIPz -HA-MAP4K4 construct was kindly shared by Vitorino and collaborators [17]. The MAP4K4 sequence was cloned into a pLVpuro-CMV-N-EGFP (Addgene, #122848) lentiviral plasmid using the gateway system. The following primers were used to mutagenize MAP4K4 to Kinase Dead (D153N) (forward: 5’-GTGATTCACCGGAACATCAAGGGCC-3’, reverse: 5’-ATGATGAATGTGAAGATGTGCCAGTCCCC-3’) and to delete the CNH domain (forward: 5’-GGTGGCAGCAGTCAGGTTTATTTCATGACCTTAGGCAGG-3’, reverse: 5’-TTTACGAATCTCCGGGGTGTCACTCTGTGGCCTAGT-3’). The MAP4K4 constructs were co-transfected with pCMV-VSVG and psPAX2 in HEK-293T to generate lentiviruses, using PEI. A431 or MDCK cells were infected with the virus particles using polybrene and selected using puromycin (0.5μg/mL). HA-MAP4K4 was also cloned into a pEGFP_C1 backbone plasmid using the restriction enzymes KpnI and HpaI. This plasmid was used for transient transfection for time-lapse imaging of MAP4K4 localization.

pLenti.PGK.Lifeact-GFP.W construct (Addgene, #51010) was co-transfected with pCMV-VSVG and psPAX2 in HEK-293T to generate lentiviruses. A431 cells were infected to stably express the Lifeact-GFP marker. Those cells were used for the migration assay and morphodynamical analysis (Fig. 1).

Paxillin-pEGFP (Addgene, #15233) construct was cloned into a pLVpuro-CMV-N-EGFP (Addgene, #122848) lentivirus plasmid using the gateway system (ATTB sites were added at the Paxillin-peGFP plasmid using the primers: forward: 5’-GGGGACAAGTTTGTACAAAAAAGCAGGCTTCGACGACCTCGACGCCCTGCTG-3’, and reverse: 5’-GGGGACCACTTTGTACAAGAAAGCTGGGTCCTAGCAGAAGAGCTTGAGGAAGC-3’), co-transfected with pCMV-VSVG and psPAX2 into HEK-293T to generate lentiviral particles and A431 were infected as mentioned before.

Lentiviral expression plasmids for E-Cadherin-mRuby3 and P-Cadherin-mCitrine were kindly shared by Arnold Hayer (McGill University, Montreal). Lentivirus particles were produced by co-transfection of pCMV-VSVG, pMDLg and pRSV-rev into HEK-293T cells and A431 cells were infected and selected with puromycin (0.5μg/mL) to stably express the cadherins as cell-cell junction markers. Those cells were used for the traction forces microscopy assay (Fig. 4).

MAP4K4 KO cells were generated using the pLenti.Cas9-blast (Addgene, #52962) construct and the following sgRNA constructs: MAP4K4_sg1 (Addgene, #76263) and MAP4K4_sg2 (Addgene, #76264). Control cells were generated using pLenti.Cas9-blast (Addgene, #52962) and the non-targeting control gRNA (Addgene, #80263). Lentivirus were produced in HEK-293T using the pCMV-VSVG and psPAX2 plasmids. Cells were co-infected with the Cas9 and sgRNA lentiviruses and co-selected with puromycin (0.5μg/mL) and blasticidin (1μg/mL).

pLVpuro-CMV-N-EGFP-MAP4K4 resistant to the MAP4K4_sg2 sequence was generated by introducing silent mutations into the targeted sequence. The following primers were used: forward: 5’-GTCAGCGCTCAGCTGGACAGGACTGTG-3’, reverse: 5’-GCCGAAATCCACAAGTTTCACCTCTGCATTCTCAGTC-3’.

Plasmids containing the sgRNA for Moesin, Ezrin and Radixin, as well as the control Rosa were a gift from Sebastien Carréno (IRIC, Montréal). Those plasmids also contain the sequence for Cas9. Cells were infected and selected using puromycin (0.5μg/mL).

### Drug treatment

GNE-495 (MedChemExpress, HY-100343) was dissolved in dimethylsulfoxide (DMSO) and diluted in complete medium. Cells were treated at the doses 0.1μM, 0.5μM or 1.0μM for 24h. The inhibitors PF-06260933 (MedChemExpress, HY-19562) and DMX-5804 (MedChemExpress, HY-111754) were also dissolved in DMSO and used at the doses 0.5μM or 1.0μM for 24h. The Rho Kinase (ROCK) inhibitor (Y-27632, cell signaling) was used to inhibit contractility after 24h of GNE-495 treatment, at 2.5 μM for 15min.

### Migration assay, live imaging and morphodynamic analysis

For migration assay, a 200μL collagen I/ matrigel gel mix at a concentration of approximately 4.5 mg/ml collagen I (Corning, 354249) and 2 mg/ml Matrigel (Corning, 354234) was added to an 8-well glass-bottomed cell culture slide (IBIDI) and let to polymerize at 37°C for 1h [66, 67]. A431 stably expressing Lifeact-GFP were plated on the collagen-matrigel gel at a concentration of 10000 cells/well and cultured for 24h. Slides were transferred into live-cell imaging mount on an inverted LSM700 confocal microscope (Zeiss) to maintain 5% CO_2_ and 37°C during movie acquisitions. Time-lapse of 5min interval were acquired with a 20× Plan Apo, NA 0.8, DICII objective, using Zen software. Clusters with approximately 6-15 cells were tracked manually by using the ImageJ

[68] plugin “Manual Tracking”, and tracking was stopped when clusters merge each other. The recorded x/y position was analyzed using the chemotaxis tool from IBIDI to calculate accumulated distance and velocity (http://ibidi.com/software/chemotaxis_and_migration_tool/). Any tracking with less than 5h was excluded from accumulated distance calculation.

For periphery displacement and extension/retraction velocities calculation, A431 cells stably expressing Lifeact-GFP were plated on 4-wells glass-bottomed cell culture slide (IBIDI) and cultured for 24h. Slides were transferred into live-cell imaging mount on a Leica SP8 confocal fluorescence microscope (Leica Microsystems) to maintain 5% CO_2_ and 37°C during movie acquisitions. A z-scan of 3 planes of clusters containing 6-15 cells was done within a 10min interval time. Acquisition was made with a 40x / 1.3 PlanApo DIC, using LasX software. Images were processed on the ImageJ software using the “Sum Intensity Z-projection”. Cluster periphery detection and extension/retraction velocities were calculated using the ADAPT plugin on ImageJ, created by David J. Barry et al [69]. For periphery displacement calculation, the “velocity visualization” output containing the detected periphery for each timepoint was opened sequentially in ImageJ and merged in a stack. The stacked image was centered using the “Template Matching” plugin created by Tseng, Q. et al. [70], and projected using the “z-project” tool. By using the “straight line” tool, we measured the thickness of the cluster border in 6 different positions, which represents the movement of the cluster periphery overtime. The velocity values were directly generated by the ADAPT plugin and plotted in GraphPad Prism.

For eGFP-Paxillin or eGFP-MAP4K4 time-lapse acquisition, A431 cells stably expressing eGFP-Paxillin or transfected with eGFP-MAP4K4 were plated on a 4-wells glass-bottomed cell culture slide (IBIDI). Cells were transferred into live-cell imaging chamber mount on a Zeiss LSM880 confocal to maintain 5% CO_2_ and 37°C during movie acquisitions. Time-lapse was acquired with a ×63/1.4 Plan Apochromat oil immersion objective, using Zen software. An interval time of 30sec was used for eGFP-paxillin, while 1min interval was used for eGFP-MAP4K4.

### Protein extraction and immunoblotting analysis

Cells were rinsed in PBS and lysed in lysis buffer (1MTris-HCl pH7.4, 1%SDS) supplemented with 100mM PMSF and the protease inhibitor cocktail (Sigma, 11697498001), on ice. Cell lysates were centrifuged for 30min at 4°C and the supernatant was collected. Protein concentrations were calculated using the BCA Protein Assay Kit (Thermo Fisher Scientific). Proteins were separated using SDS-PAGE gels (8%, 10% or 12%, according to protein size) and transferred into PVDF membranes. The membranes were blocked in skim milk 5% for 1h and exposed to the primary antibodies diluted in TBS-tween 0.1%, BSA2% overnight at 4°C. Secondary antibodies were incubated at RT for 1h and membranes were revealed in an X-ray film in a dark room.

### Immunofluorescence

Cells were plated on coverslips to reach cluster confluence as described before. Cells were fixed with 4% PFA for 20min at RT. Exceptionally, for pERM staining requires fixation with 10% TCA on ice. Cells were rinsed 3x with PBS and permeabilized with 0.1% Triton X-100 for 2min. After rinsed 3x with blocking solution (2% BSA in PBS), the coverslips were incubated in blocking solution for 1h at RT. Primary antibodies were diluted in blocking solution and incubated for 1-3h at 37°C or overnight at 4°C, both in humidified chamber. Secondary antibodies were co-incubated with phalloidin and DAPI for 1h at RT, diluted in blocking solution. The slides were mounted using mowiol mounting medium or vectashield (Vector Laboratories Canada, H-1000).

### Image acquisition and processing

Images were acquired using the confocal microscopes: LSM 700 (Carl Zeiss), coupled to a ×63/1.4 Plan Apochromat DIC oil immersion objective, LSM880 (Carl Zeiss), equipped with a ×63/1.4 Plan Apochromat oil immersion objective, or a Leica SP8 (Leica Microsystems), equipped with 63x /1.4 PlanApo DIC immersion oil objective.

Cluster circularity was calculated using the “shape descriptor” measurement from ImageJ, using the “freehand selection” tool to draw the cluster as the ROI (region of interest). Protrusion areas were calculated also using the “freehand selection” tool from ImageJ. We considered as a protrusion the actin area in front of the nucleus that do not present retraction fibers, and were constrained laterally by the actin arches for DMSO, or cell-cell junctions for GNE-495 treated cells. Clusters stained with phalloidin and DAPI were used for those analysis.

For F-actin network analysis, we made a z-scan of the entire cluster stained with phalloidin and DAPI. Images were done every 0.3μm. The first plane was used to measure ventral fibers, while the dorsal actin network was done using the “maximum intensity” projection (ImageJ) from the focal plane at the middle of the nucleus to the top of the cluster. Using imageJ, we plotted the intensity profile of a line scan manually traced from the distal to the proximal part of protruding cells, considering at least 3 cells per cluster. A minimum intensity threshold was chosen for each experiment. Each intensity maxima above this threshold were considered a bundle of F-actin and counted.

Dorsal stress fibers analysis was done using a z-scan of entire clusters stained for F-actin and α-actinin, with a z interval of 0.3μm. The 3D reconstitution of the clusters was done using the Imaris software (Bitplane), and the F-actin structures enriched in α-actinin and elongating from the bottom of the cluster to its dorsal part were counted. The estimated length in 3D was also measured manually using Imaris (Bitplane).

Mature focal adhesions number was calculated using the Surface tool on Imaris (Bitplane), from clusters stained for zyxin and F-actin. A minimum threshold of 3μm^2^ surface was set to also select the mature focal adhesions by size.

Mean Intensity of pERM staining was measured using ImageJ by using the “freehand selection” tool to draw the cluster as the ROI. Number and length of retraction fibers were calculated from clusters stained for pERM. We measured the length manually, by using the “segmented line” tool on ImageJ. Background intensity was systematically subtracted.

Intensity of pMLC2 was measured using a z-scan of entire clusters, with a z interval of 0.3μm. “Sum intensity” projection was done on ImageJ. Cluster and protrusion ROI were selected as mentioned before, using the phalloidin and DAPI channels. The mean intensity and the area of each ROIs were measured and multiplied to calculate the total intensity. Background intensity was systematically subtracted.

Cell-cell junction tortuosity was measured using the “maximum intensity” projection of z-scan of entire clusters stained for p120 catenin. Images were done every 0.3μm. The junction signal at one side of a cell-cell junction was outlined using the “segmented line tool” on Image J. Then, we measured the length between the two extremities of this outline using a straight line. The tortuosity was calculated using the ratio of the outline length over the straight length.

For vinculin accumulation at the cell-cell junction, we used the p120 channel to select the ROI and vinculin colocalization was calculated using the ImageJ “colocalization test” tool, selecting the Pearson’s colocalization output.

Line scans for measuring MAP4K4 localization at cell-cell junctions were done using the “Sum intensity” projection tool from ImageJ, with a z-scan containing the high of the cell-cell junction. Images were taken every 0.7μm.

### Cell scattering assay

A431 cells were infected with pLVpuro-CMV-N-EGFP-MAP4K4 (WT, kinase dead or deleted for the CNH domain) and plated on coverslips to induce cluster formation, as described before. Cells were immunostained for F-actin and nuclei. Using the LSM 880 (Carl Zeiss), coupled to a 20x / 0.8 PlanApo DIC, we acquired tiles of 2×2 in 8 different regions of the coverslip, randomly selected. The total number of cells per field was determined using the Spot tool on Imaris (Bitplane) to detect the nuclei. The mean eGFP intensity was calculated using the Surface tool on Imaris (Bitplane) on the F-actin channel to detect the total cell area as the ROI. The total eGFP intensity was calculated by multiplying the mean intensity by the cells’ area. Background intensity was systematically subtracted.

### Synthesis of PDMS silicone substrates

Compliant polydimethylsiloxane (PDMS) substrates of known stiffness were prepared as described previously [71, 72]. To summarize the manufacture, parts A and B of NuSil 8100 (NuSil Silicone Technologies, United States) were mixed at a 1:1 w/w ratio. The stiffness of the silicone substrates was then tuned by adding a certain concentration of Sylgard 184 PDMS cross-linking agent (dimethyl, methyl hydrogen siloxane, containing methylterminated silicon hydride units) to the PDMS. The mechanical properties of these PDMS substrates have previously been extensively characterized [71, 72]. For our experiments, we selected a concentration of 0.36% w/w Sylgard 184 crosslinker, resulting in a 12kPa stiffness as this is within the range of *in vivo* stiffness epidermoid carcinoma and resembles a stiffer tumour microenvironment. 50μl of uncured PDMS and spread onto square ∼ 22 mm (no. 1) glass coverslips, then cured for 1h at 100ºC to yield silicone substrates with a 100μm thickness. DID-conjugated (far-red) fluorescent fiduciary beads were synthesized as described previously [73], mixed into uncured PDMS and crosslinker, then spin-coated onto the PDMS substrates at 3000rpm for 1 minute to yield a bead-embedded PDMS layer ∼ 1 μm thick (WS-650 Spin Processor, Laurell Technologies). The substrates were then incubated at 100ºC for 1h. They were then fastened to the bottom of 6-well plates, functionalized using Sulfo-SANPAH, and then protein coated with collagen for cell adhesion.

### Traction force microscopy and Bayesian inversion stress microscopy

Cell-generated surface displacements, traction stress, and strain energy were quantified using traction force microscopy (TFM) as described in literature [71, 74] using an open-source Python TFM package (pyTFM) modified to the case of cell clusters and force imbalances within the field of view [36, 75, 76]. Intercellular stresses, shear and normal stresses were then quantified from the traction forces using Bayesian inversion stress microscopy (BISM) using a MATLAB package [77]. Cells were plated, allowed to settle and form clusters 48h prior to imaging. Prior to imaging, the cells were treated with HOECHST 33342 Dye (Thermo Fisher Scientific, Inc.) in order to stain the nuclei. During imaging, A431 cells expressing mCitrine-tagged P-cadherins and A431 cells expressing mRuby-tagged E-cadherins were used. We focused on imaging isolated cell clusters in a clear field of view consisting of 5-13 cells. For imaging, the 6-well plate containing the cells were mounted onto a confocal microscope (Leica TCS SP8 with a 10Å∼0.4 NA objective). They were then maintained at 37ºC (stage heater, Cell MicroControls, VA) and 5% CO_2_ (perfusing 100% humidity pre-bottled 5% CO_2_ in synthetic air). The cells, nuclei and fiduciary TFM beads were simultaneously imaged using fluorescent and transmission microscopy over several hour time courses at time intervals of 15-40 min. The resulting images were then corrected for lateral drift using an ImageJ pipeline, the values outside the cell cluster were masked to remove the background noise, then analyzed using the pyTFM and BISM workflows to obtain the displacements, tractions, strain and intercellular stresses.

### Statistical analysis

All graphs and statistical tests were done using GraphPad Prism (GraphPad Software). We performed at least 3 independent experiments (N) for each analysis and the minimum number of data points (n) is specified at figure legends. Since normal distribution of the data was not formally tested, we used the non-parametric Mann-Whitney test or Kruskal-Wallis test with Dunn’s correction for multiple comparisons of unpaired data. Paired data was analyzed using Wilcoxon test. Values are expressed as mean±s.d., unless otherwise indicated at the figure legend, and all individual values are represented at the graphics. *P*-values are noted on figures as following: \**P*<0.05, \*\**P*<0.01, \*\*\**P*<0.001, \*\*\*\**P*<0.0001.

## ACKNOWLEDGMENTS

We thank Arnold Hayer (McGill U.), Sebastien Carréno (U. Montreal) and Weilan Ye (Genentech) for their generosity in sharing reagents. We thank C. Charbonneau for technical assistance and the entire Emery lab and Arnold Hayer for helpful discussions and critical reading of the manuscript. This work was supported by grants from the Canadian Institute for Health Research (CIHR; PJT – 175093 to G.E. and A.J.E., and #143327 to A.J.E.), from the Natural Sciences and Engineering Research Council of Canada (NSERC, RGPIN-2020-07169 to A.J.E) and from Canadian Foundation for Innovation (Project #32749 to A.J.E.). L.E.A.D held a doctoral scholarship and C.P. a postdoctoral fellowship from Fonds de Recherche du Québec – Santé (FRQS).

---

**Extended Data, Fig. 1.**
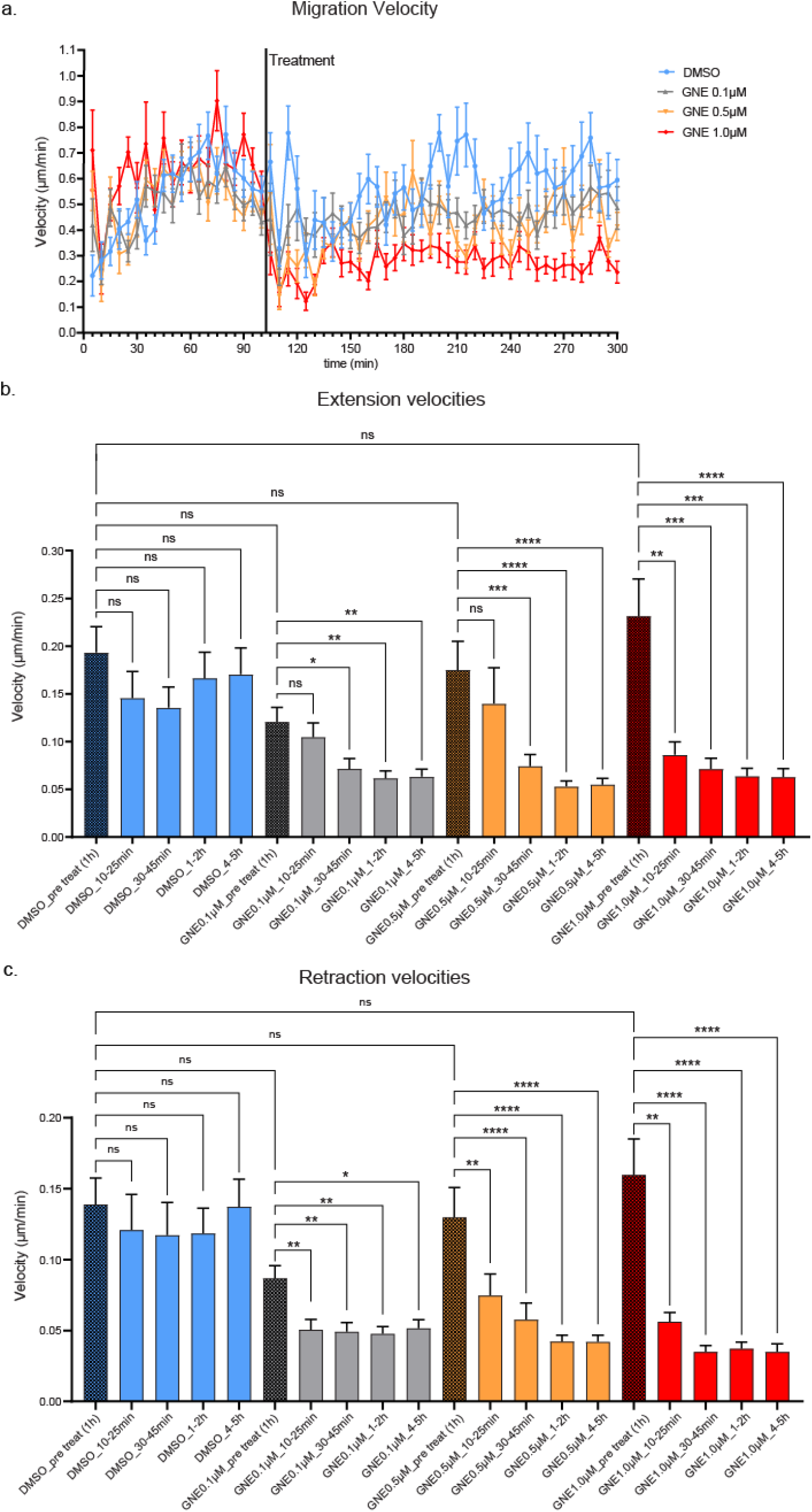
MAP4K4 inhibition decreases cluster migration in a dose-time dependent way. (a) mean migration velocities of A431 clusters at each acquired time point, before and after the treatment with DMSO or indicated doses of GNE-495. (b, c) Mean extension (b) and retraction (c) velocities of A431 clusters, treated with DMSO or indicated GNE-495 doses, represented as a mean of the indicated periods of time. Data on (a) is represented by mean±sem. Data on (b) and (c) are represented as mean±s.d. and tested by Kruskal-Wallis (ns: nonsignificant, \**p*<0.05, \*\**p*<0.01, \*\*\**p*<0.001, \*\*\*\**p*<0.0001).

**Extended Data, Fig. 2.**
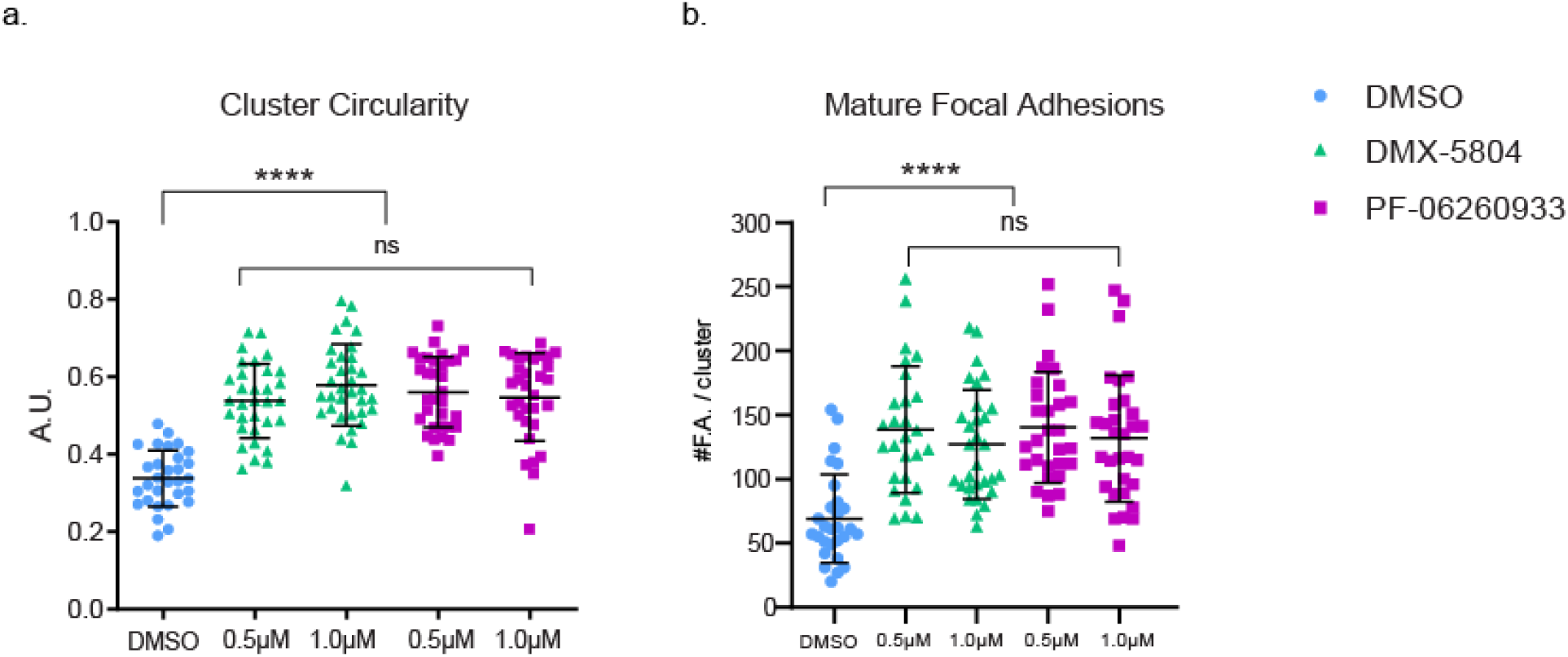
Effect of MAP4K4 inhibition with different specific inhibitors. (a) circularity of clusters treated with DMSO, DMX-5804 or PF-06260933 compounds, at the indicated doses. At least 28 clusters from 3 independent experiments were analyzed. (b) number of zyxin-positive focal adhesions on clusters treated with DMSO, DMX-5804 or PF-06260933 compounds, at the indicated doses. At least 25 clusters from 3 independent experiments were analyzed. Data are represented as mean±s.d. and tested by Mann-Whitney (ns: nonsignificant, \**p*<0.05, \*\**p*<0.01, \*\*\**p*<0.001, \*\*\*\**p*<0.0001).

**Extended Data, Fig. 3.**
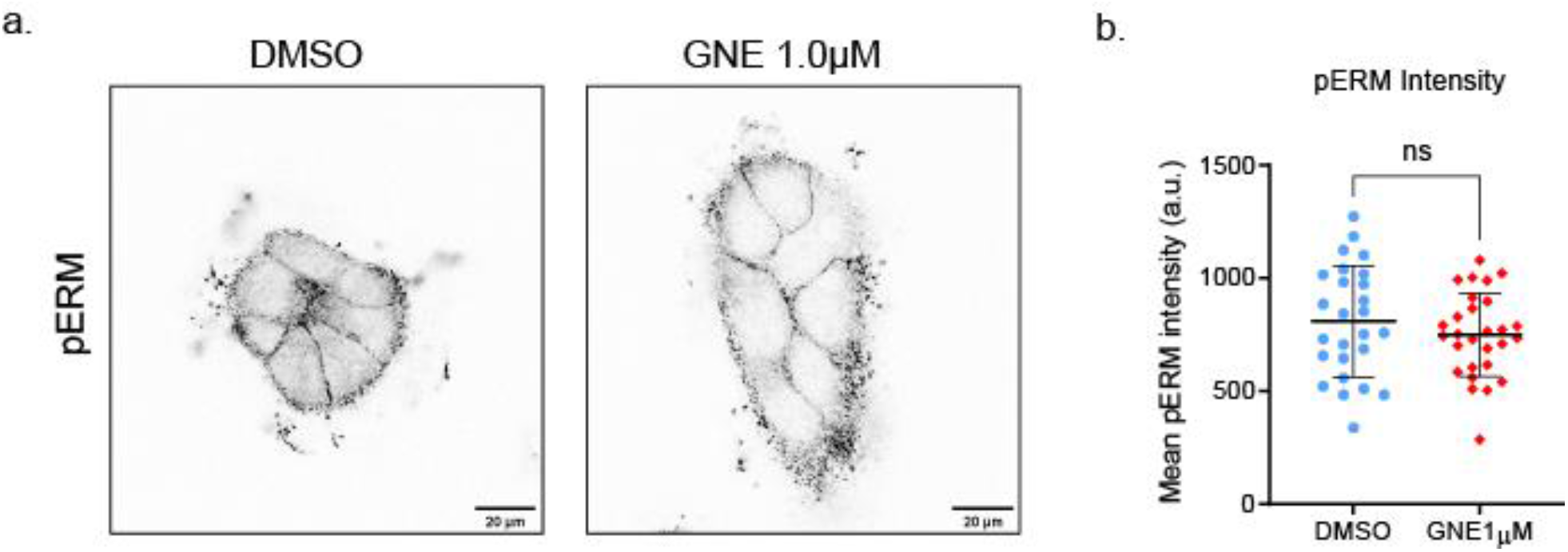
MAP4K4 does not regulate junction tortuosity through Moesin phosphorylation. (a) representative confocal images of pERM staining at the cell-cell junction z focal plane, from clusters treated with DMSO or GNE-495 at 1.0μM. (b) quantification of mean intensity of pERM of clusters treated with DMSO or GNE-495 at 1.0μM. At least 25 clusters from 3 independent experiments were analyzed. Data are represented as mean±s.d. and tested by Mann-Whitney (ns: nonsignificant, \**p*<0.05, \*\**p*<0.01, \*\*\**p*<0.001, \*\*\*\**p*<0.0001).

**Extended Data, Fig. 4.**
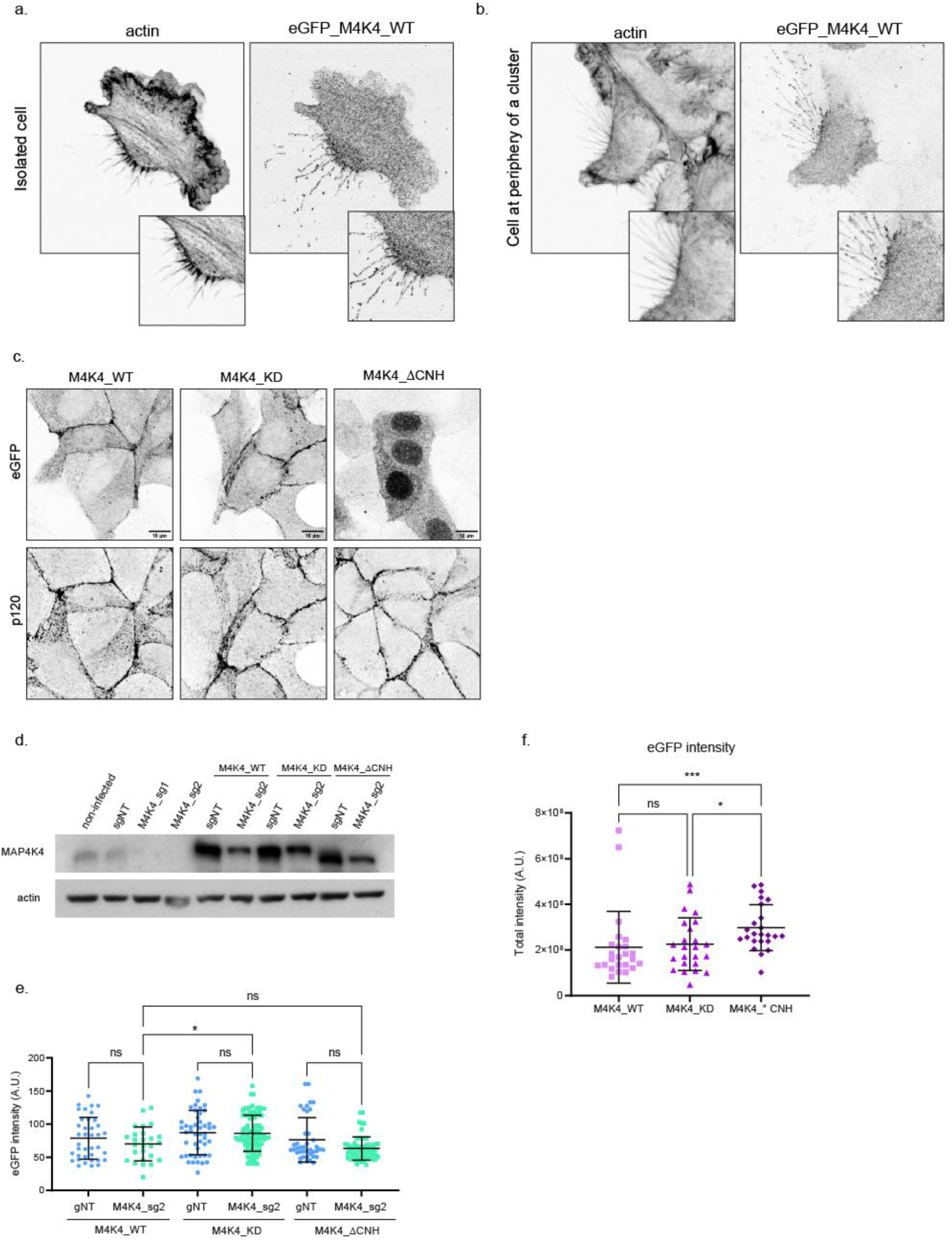
MAP4K4 localizes at retraction fibers and adherens junctions in A431 cells and MDCK cells. (a, b) representative confocal images of an isolated cell (a) or cell at cluster periphery (c) retracting, expressing eGFP-MAP4K4_WT, showing MAP4K4 localization at retraction fibers and at the base of the retraction in A431 cells. (c) representative confocal images of confluent MDCK cells expressing eGFP-MAP4K4_WT, KD or ΔCNH. Staining of p120-catenin indicates cell-cell junctions. (d) Immunobloting of MAP4K4 or actin for lysates of A431 cells controls (non-infected or sgNT), KO for MAP4K4 (sg_1 or sg_2) alone or expressing eGFP-MAP4K4 WT, KD or ΔCNH, resistant to sg_2. (e) eGFP mean intensity of cells control or KO for MAP4K4 (sg2), stably expressing eGFP-MAP4K4 WT, KD or ΔCNH, resistant to sg_2, considered on the tortuosity index quantification (Fig. 6f). (f) eGFP total intensity of cells stably expressing eGFP-MAP4K4 WT, KD or ΔCNH, considered to calculate % of cell isolation (Fig. 6h).

**Video 1. MAP4K4 inhibition decrease focal adhesion dynamics**. Confocal time-lapse recording of focal adhesion dynamics on DMSO (left) or GNE-495 at 1.0μM (right) treated A431 cells. Cells stably expressing Paxillin-eGFP, shown in grayscale. Time-interval: 30sec.

**Video 2. MAP4K4 localize at retraction fibers and cell rear, as well as cell-cell contact**. Confocal time-lapse recording of A431 cells expressing MAP4K4-eGFP (WT), shown in grayscale (left panel) or overexposed (right panel). Yellow arrowheads indicate accumulation of MAP4K4 at cell rear and retraction fibers. Red arrows indicate cell-cell contacts. Cyan arrows indicate accumulation of MAP4K4 at cell-cell contacts. Time-interval: 1min.

## Notes

### Competing Interest Statement

The authors have declared no competing interest.

